# Impact of blood storage conditions on the transcript profile of plasma cell-free RNA

**DOI:** 10.1101/2021.03.30.437637

**Authors:** Jinghua Sun, Xi Yang, Taifu Wang, Yanru Xing, Haixiao Chen, Sujun Zhu, Juan Zeng, Qing Zhou, Fang Chen, Xiuqing Zhang, Wen-Jing Wang

**Affiliations:** BGI Education Center, University of Chinese Academy of Sciences, Shenzhen 518083, China; BGI-Shenzhen, Shenzhen 518083, China; Obstetrics Department, Shenzhen Maternity and Child Healthcare Hospital, Shenzhen, Guangdong Province, China; Shenzhen Engineering Laboratory for Birth Defects Screening, BGI-Shenzhen, Shenzhen 518083, China

**Keywords:** plasma cell-free RNA, blood storage, RNA contamination, Transcriptome

## Abstract

**BACKGROUND:** Plasma cell-free RNA (cfRNA) are potential biomarkers for disease prediction and diagnosis. However, pre-analysis factors, such as the delay in blood processing and storage may lead to unreliable results, though no study has systematically evaluated the effect of blood storage conditions on the whole transcriptome of plasma cfRNA yet.

**METHODS:** We collected peripheral blood samples from four healthy subjects and allowed them to stand at room temperature or 4◻ for different time periods (0h, 2h, 6h and 24h) prior to plasma separation. Then, plasma cfRNA stability was evaluated by measuring expression changes of cell-free mRNA, lncRNA and miRNA using high throughput sequencing-based profiling. Finally, their paired leukocyte RNA data were integrated to depict the effect of leukocytes on plasma cfRNA during storage.

**RESULTS:** Plasma mRNA and lncRNA presented high correlations (Pearson R^2^ ≥ 0.8) and fewer variations when blood was stored at 4◻ for 6 hours or stored at RT for 2 hours. miRNA was more stable, with minimal R^2^ of 0.86 at 4◻ for at least 24 hours or at RT for 6 hours. Correlations of plasma RNA and leukocyte RNA increased with the incubation time, and the relative proportion of neutrophils in plasma grown from 14.3% to 61.2% at RT (*P* = 0.004), indicating leukocyte RNA contamination. Besides, the tissue enriched genes in plasma were down-regulated with the extension of storage time.

**CONCLUSIONS:** Our results characterized the effects of short-term storage of blood samples on plasma cfRNA, which will facilitate further researches or clinical applications to avoid bias resulting from sample processing.

## Introduction

In the past few years, an increasing number of studies have reported the potential use of plasma cell-free miRNAs (cf-miRNA), lncRNA (cf-lncRNA) and mRNA (cf-mRNA) as circulating biomarkers for monitoring, prediction or diagnosis of a wide variety of diseases, such as cancers [1], pregnancy complexities [2, 3] and Alzheimer‘s disease [4]. However, several limitations of cfRNA-based liquid biopsies have been addressed in the past few years [5], including pre-analytical conditions, which may lead to discordant results. Pre-analytical challenges that include the short half-life of cfRNA as well as cellular RNA contamination, standardization of the sample process workflow will benefit in reducing these variations in the clinical implementation of plasma cfRNA.

In real-life clinical practice, short-term storage or transportation of blood samples from the collection site to the laboratory is commonly required during research or clinical application, such as multicenter clinical trials. In order to achieve consistent results, it requires evaluating the impact of short-term storage of blood on the expression of cfRNA. A limited number of studies have been performed to determine optimal blood storage conditions for plasma cfRNA analysis. However, all of these studies only focused on specific genes, such as cancer-associated [6], placental-specific [7], or established housekeeping genes [8]. Yet no study has systematically evaluated the influence of blood storage conditions on the full transcriptome of cfRNA in plasma. With the development of sequencing technology of plasma cfRNA [9, 10], more informative cfRNA can be detected, which makes it possible to obtain the whole transcriptome profile of plasma cfRNA to evaluate the impact of the delay of blood processing.

Therefore, we investigated the effects of pre-analytical factors, including blood storage times (0h, 2h, 6h and 24h) and temperatures [room temperature (RT) and 4◻] on the cfRNA transcript profiles in plasma. Besides, we tried to depict the possible leukocyte RNA contamination and the decay of plasma original cfRNA through integrative analysis with paired leukocyte RNA data.

## Methods

### Sample collection and study design

This study was approved by the Institutional Review Board of Shenzhen Maternity and Child Healthcare Hospital (SFYLS 2019 NO. 139) and all participants were provided with written informed consent prior to blood collection. Fresh blood samples were obtained from four healthy subjects. The blood samples were collected in ethylenediaminetetraacetic acid (EDTA) Vacutainer tube (BD, 0202992058) and stored at RT (18-22◻) or 4◻ for various time intervals (0h, 2h, 6h and 24h) before further processing (Fig.1). At each condition, 1mL blood was taken and 300ul plasma was separated according to the two-steps centrifugation protocol: 1600g for 10 min at 4◻ and 12,000g for 10 min at 4◻. Then, 900uL TRIZOL LS Reagent (Thermo Fisher, 10296028) was added and mixed immediately to extract the total RNA according to the manufacturer’s protocol.

**Fig. 1.**
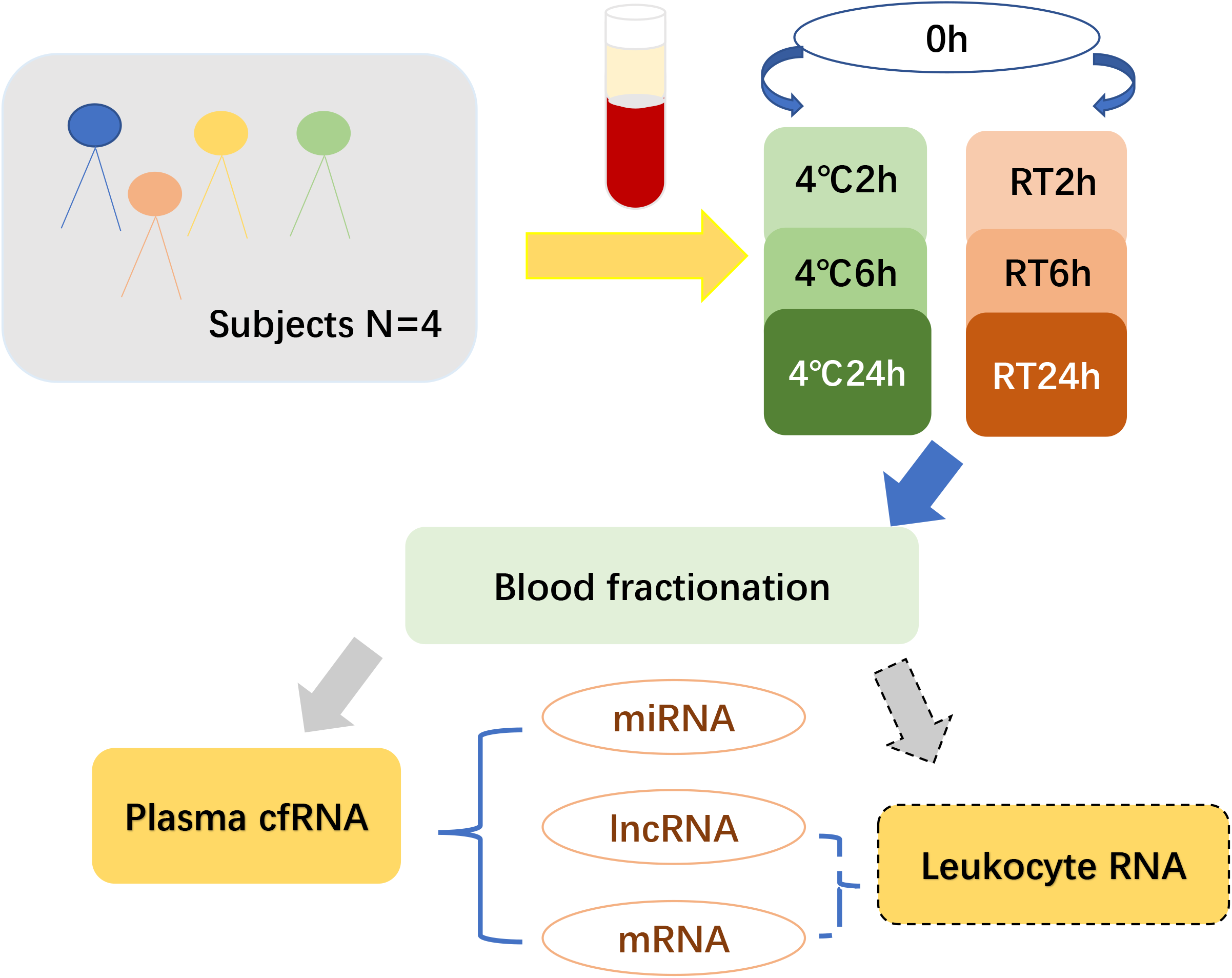
Study design. RT, room temperature.

The transcriptome library was constructed using PolyAdenylation Ligation Mediated-Seq (PALM-Seq) method [9], which is a simultaneous sequencing method for cell-free coding and non-coding RNA. cfRNA sequencing was performed on BGISEQ-500RS (single-end 100 bp) platform with an average depth of 100 million reads per sample. Erythrocyte lysis was performed in the remaining portion of the blood and the leukocyte transcriptome library was constructed with RNase H method. Details of the leukocyte RNA sample preparations were published previously by Xing et al [11]. In this study, we focused on the expression profile changes of plasma cfRNA and the possible effects of leukocytes on cfRNA.

### Alignment and Quantification

Details of cfRNA sequencing data preprocess were carried out as previously described [9]. In brief, adapter and low-quality reads were trimmed, then reads with > 10% ‘N’ base or shorter than 17bp were filtered by cutadapt [12]. The remaining reads were aligned to the hg19 by bowtie [13] with the order of rRNA and Y_RNA, miRNA, mRNA and lncRNA. The reads count of miRNA was normalized to reads per million mapped reads (RPM) using custom script, and gene-level expression of mRNA and lncRNA was estimated by RSEM [14] and was normalized to transcripts per kilobase million mapped reads (TPM). The transcript relative coverage across all transcripts was produced by RSeQC [15] with geneBody_coverage.py, in which transcripts of mRNA and lncRNA were normalized to 100 percentiles and transcripts of miRNA were normalized to 40 percentiles.

For leukocytes RNA, clean reads were obtained after rRNA filtering (using SOAP2 [16]) and the low-quality reads and adaptor filtering (using SOAPnuke [17]). Then, the clean reads were aligned to the mRNA and lncRNA transcriptome by bowtie2 [18] and quantified at gene-level by RSEM [14].

### Gene differential expression and functional analysis

Differential expression gene (DEG) analyses were performed between 0h and the other storage conditions. The genes were considered as differentially expressed across different groups if fold change (FC) ≥ 2 and adjusted p-value < 0.05 using the DESeq2 R package [19]. Expressions of specific genes were considered altered moderately if raw *p* < 0.05 in the DEG analysis. The Short Time-series Expression Miner (STEM) [20] program was used to cluster and visualize the possible profiles and changes in expression over time in DEGs. The pathway and functional enrichment analysis were performed in Metascape [21] (http://metascape.org/gp/index.html#/main/step1) with express analysis mode. Only a pathway with q-value < 0.01 considered as significant enrichment.

### Platelet specific genes and tissue enriched genes (TEGs)

Platelet-specific genes were downloaded from the PanglaoDB database [22] (https://panglaodb.se/). The genes with ubiquitousness index = 0 and marker count < 4 keep to further analysis. Tissue-specific genes were downloaded from the human protein atlas database (HPA) [23] (https://www.proteinatlas.org/). The genes with expression level greater than 1 and have at least five-fold higher expression levels in a particular tissue compared to all other tissues were classified as the TEGs. Only tissue with at least 10 TEGs will be a candidate tissue and a total of 23 tissues were included. All tissue-specific genes must have a TPM > 0 in at least 2 plasma samples and TPM ≥ 1 in at least one plasma sample were included in the TEGs analysis.

### Deconvolution analyses of plasma cfRNA

Leukocyte’s relative proportion of plasma cfRNA was estimated by CIBERSORT [24] with 100 permutations and quantile normalization disabled. The same type of cells in the resting state and activated state were merged, for example, resting CD4 memory T cells and activated CD4 memory T cells are merged into CD4 memory T cells.

### Permutation test for overlap DEG of leukocyte RNA and plasma cfRNA

The expected overlap DEG numbers of leukocyte RNA and plasma cfRNA were produced by randomly selecting genes with the same number of DEG from two groups expressed genes and calculated the overlap genes number. This step was conducted 999 times for each task, then p-values were calculated using the Monte-Carlo method with one-sided test [25].

### Statistical analysis

Statistics were performed using R (3.5.1). Unless otherwise stated, analyses were performed using two-tailed paired t-test (for matched samples) and p-values < 0.05 were considered significant. The correlations between two samples were performed using the R function lm () after the log2 transformation of gene expression. Principal component analysis (PCA) plots were generated using the function prcomp (). Venn diagrams were generated by Venny [26] (https://bioinfogp.cnb.csic.es/tools/venny/). The ratio of DEGs were calculated by the number of differential genes divided by the number of expressed genes (TPM or RPM > 0).

### Data availability

The data that support the findings of this study have been deposited into CNGB Sequence Archive (CNSA) [27] of China National GeneBank DataBase (CNGBdb) [28] with accession number CNP0001320.

## Results

In total, we generated 28 cell-free transcriptomes by high throughput sequencing method to determine the effect of different storage conditions (Fig.1). On average, each plasma sample had 64 million clean reads after removal of rRNA and Y_RNA, which led us to identify about 4804 lncRNA genes, 13701 mRNA genes and 818 miRNA per sample. To test if short-term storage affects transcript coverage, we first compared the variation in gene body coverage across all transcripts from 5′ to 3′. Among the three types of RNA, lncRNA and miRNA were highly concordant for all conditions, while mRNA had slight fluctuation at the 5’ ends (Supplementary Fig. 1), suggesting a little effect on transcript overall coverage.

### Effect of blood storage on plasma cfRNA whole transcriptome profile

To investigate the effects of storage conditions on the whole transcriptome profile, we performed a PCA analysis on all the samples. The mRNA and lncRNA were stored at RT or 4◻ for 24h were clearly distinguishable from other conditions (Fig. 2A). For miRNA, only samples stored at RT for 24h could be differentiated from other samples (Fig. 2A). To further confirm this result, we calculated the Pearson’s correlation coefficients and performed DEG analysis between 0h and the other conditions. When blood was stored at 4◻, the plasma miRNA was stable until 24h with a minimum R square of 0.86, while mRNA and lncRNA had decreasing correlations (Fig. 2B, Supplementary Table 2). DEG analysis also showed that there was no DEG at 2h and 6h, and were only 4 miRNA DEGs when blood was stored for 24h (Fig. 2C, Supplementary Table 1), revealing that short-term storage conditions slightly affect cf-miRNA at 4◻ until 24h. The effects on mRNA and lncRNA were noted when blood stored for 24h at 4◻ as the numbers of DEG increased to 234 (234/16855=1.39%) for mRNA, and 43 (43/8567=0.5%) for lncRNA at 24h (Fig. 2B), which is in line with PCA result, indicating cf-mRNA and cf-lncRNA are relatively stable for 6h but not 24h at 4◻. Compared with 4◻, blood stored at RT had lower correlations and more DEG at the same time point, especially at 24h. When blood was stored at RT for 24h, the mean R^2^ of mRNA and lncRNA decreased to 0.63 and 0.71 (Supplementary Table 2), with 3281 (3281/16792=19.54%) and 1100 (1100/8145=13.5%) DEGs, respectively (Fig. 2C). Besides, we noticed that the average correlations of mRNA and lncRNA at RT for 6h are smaller than that at 4◻ for 24h (mRNA: 0.73 vs. 0.76; lncRNA: 0.78 vs. 0.80) (Fig. 2B, Supplementary Table 2), implying a greater effect on mRNA and lncRNA when blood was stored at RT for 6h. miRNA kept high correlations (R2 > 0.85) at RT during storage, however, the expression of 54 (54/1360=3.97%) miRNA genes changed significantly at 24h (Fig. 2B, 2C Supplementary Table 2).

**Fig. 2.**
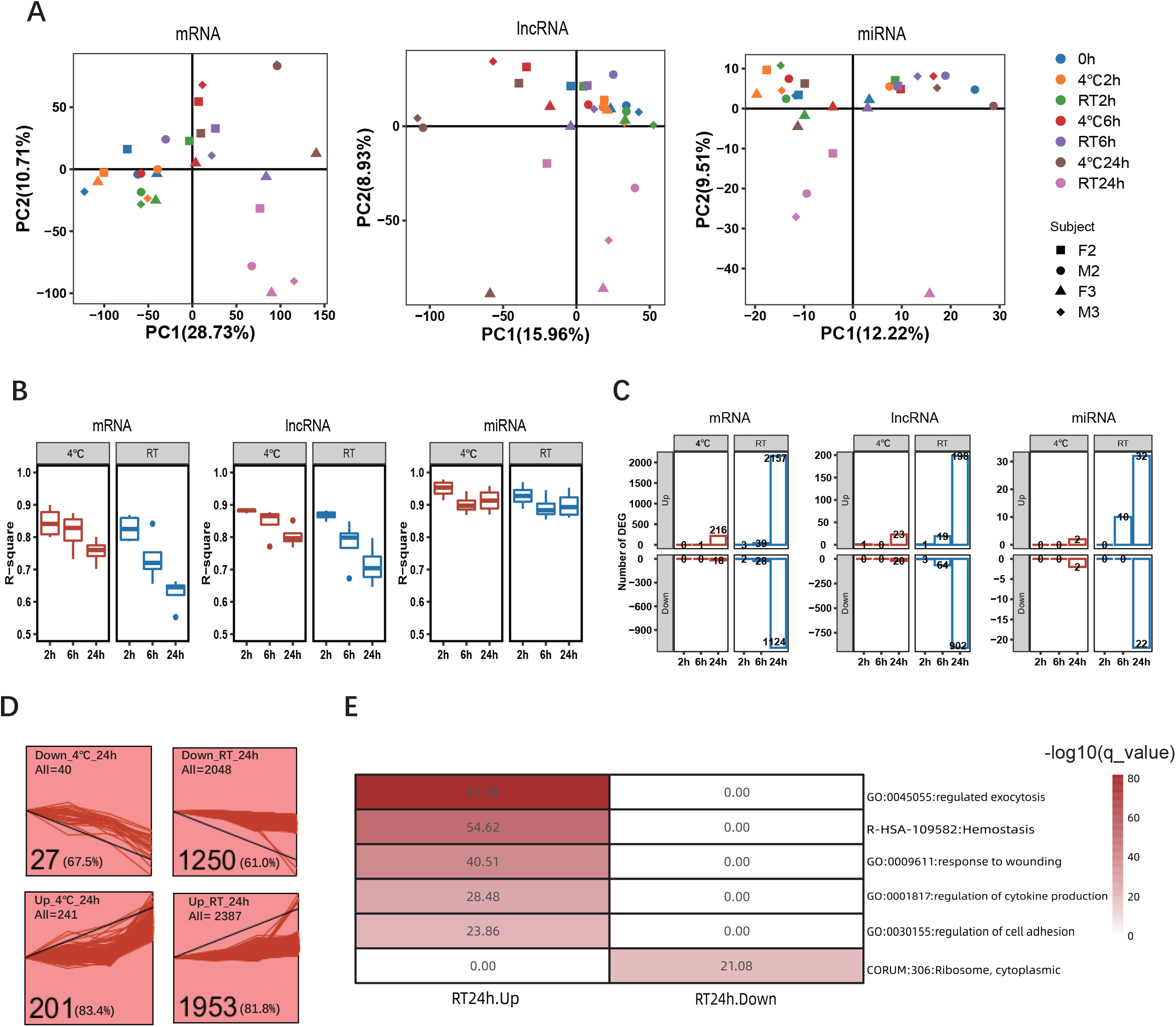
Effect of blood storage on plasma cfRNA overall transcriptome profile. (A) PCA plot on samples with different storage conditions. (B) Correlations of gene expression between 0h and the other conditions. R^2^ is the square of Pearson’s correlation coefficient. (C) Different expression gene numbers at each condition, compared with 0h. (D) time-series cluster profile for all DEGs. (E) Enriched pathway of significantly up-regulated and down-regulated genes at RT after 24h incubation. DEG, Differential expression gene.

To identify significantly represented temporal gene expression profiles during storage, we further performed time series analysis on DEGs at 24h. As a result, the two most significant profiles corresponded to gradual decrease (p < 0.05) and increase (p < 0.05) in expression, accounting for about 80% up-regulated genes and more than 60% down-regulated genes, regardless of blood storage temperature (Fig. 2D), which implies that the influence of storage conditions on the plasma expression profile is a process that gradually increases over time.

To reveal the biological processes that may occur during blood storage, we conducted gene functional enrichment analysis on mRNA DEGs. Only DEGs of 24h at RT was found to be enriched in some pathways or protein complexes. On the one hand, the up-regulated genes of 24h at RT are mainly involved in the biological processes of regulated exocytosis, hemostasis, response to wounding, regulation of cytokine production and regulation of cell adhesion (Fig. 2E). Among these, we noticed that leukocyte activation and degranulation related processes are the members of exocytosis and regulation of cell adhesion pathways (Supplementary Table 3). Besides, we also observed an increased expression of 8/10 platelet-specific genes at RT for 24h, while no gene was up-regulated significantly at leukocyte, although most genes had an increasing trend (Supplementary Fig. 2) at leukocyte. On the other hand, we found down-regulated genes of 24h at RT are enriched in cytoplasmic ribosomal protein complex (Fig. 2E, Supplementary Table 3).

### Effect of blood storage on established housekeeping genes

Housekeeping genes are thought to be a stable expression and therefore are widely used as internal controls for reverse transcription polymerase chain reaction (RT-qPCR). To explore whether these established housekeeping genes are stable under different blood storage conditions, eighteen housekeeping genes [29] for mRNA and six housekeeping genes [30] for miRNA were examined. For long RNA, when blood was stored at 4◻, no housekeeping gene was significantly altered, only GUSB (*p* = 0.018) altered moderately and several genes such as PSMB4, SNRPD3 and TFRC were little impacted over time (Fig. 3A). However, when blood was stored at RT, 4/18 genes were significantly up-regulated at 24h, including GUSB, ACTB, RAB7A and HSP90AA1 (Fig. 3A). Totally, 5/18 genes were moderately up-regulated at 24h while SNRPD3 was moderately down-regulated. Meanwhile, CHMP2A, TUBB, C1orf43, VPS29 and PSMB2 were relatively stable at both 4◻ and RT with time (Fig. 3A), which can be candidate reference genes for RT-qPCR. For housekeeping genes of miRNA, only hsa-miR-106a-5p significantly decreased at RT after 24h compared to 0h (Fig. 3B). The other genes, such as hsa-miR-93-5p and hsa-miR-17-5p were moderately down-regulated after 24h incubation of whole blood at RT (Fig. 3B). hsa-miR-16-5p, hsa-miR-191-5p and hsa-miR-25-3p were relatively stable at both 4◻ and RT over time (Fig. 3B), which can be candidate reference genes for miRNA RT-qPCR.

**Fig. 3.**
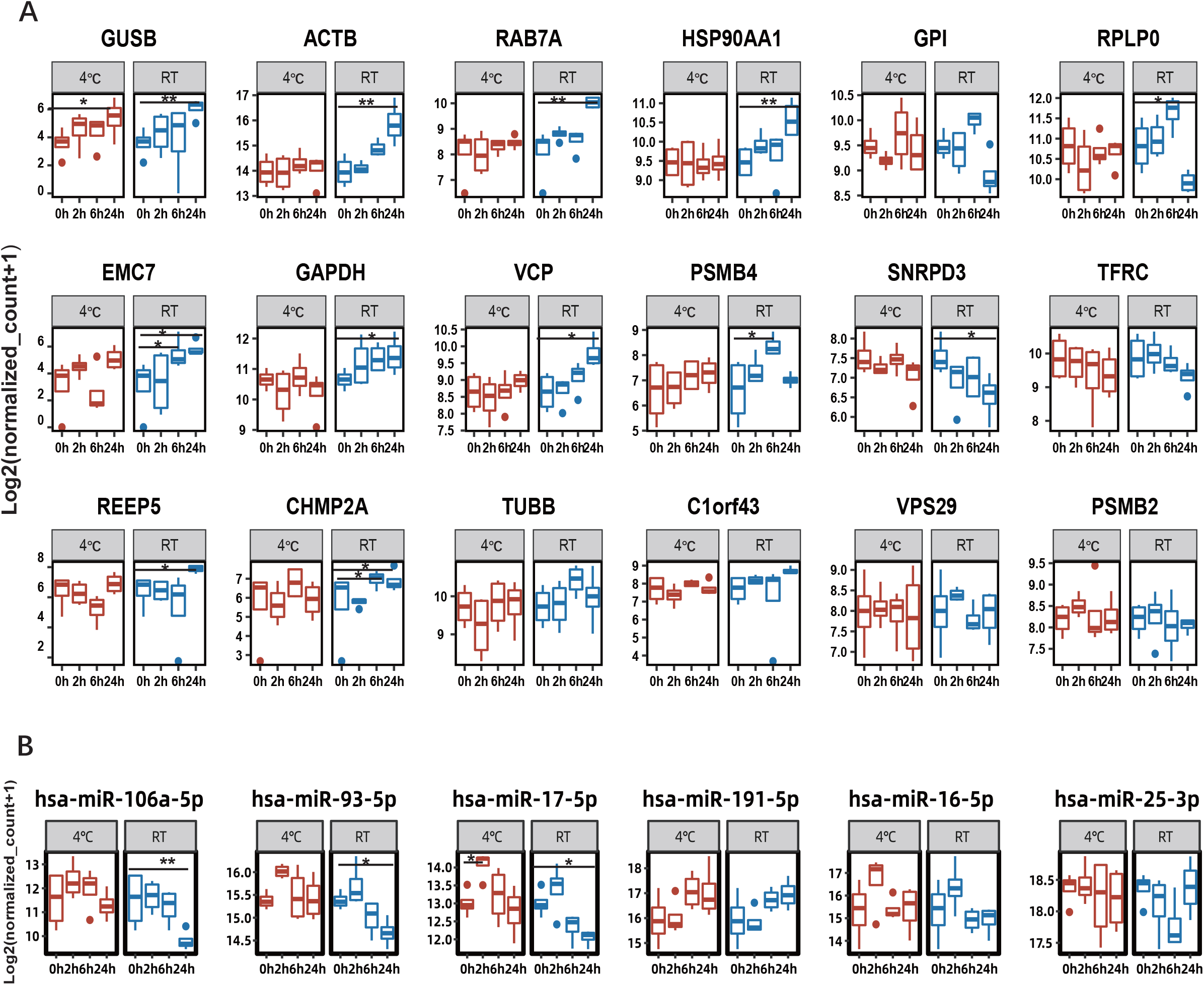
Effect of blood storage on established housekeeping genes. (A) Dynamic expressions of mRNA housekeeping gene during blood storage; (B) Dynamic expressions of miRNA housekeeping genes during blood storage. The raw counts were normalized by DESeq2. ** differential expression genes, that is log2 (fold change) ≥ 1 and adjusted p-value<0.05; * genes with moderately changed expression, that is only *p* < 0.05.

### Effect of leukocyte on plasma cfRNA during blood storage

For plasma cfRNA, leukocyte RNA contamination is a great challenge for the delay of blood processing. To evaluate the contamination of leukocyte RNA, we collected corresponding leukocyte RNA data from a previous study. Increasing correlations of cfRNA and leukocyte RNA profile were observed over time, both at RT or at 4◻ (Fig. 4A), suggesting the leukocyte RNA contamination is increasing.

**Fig. 4.**
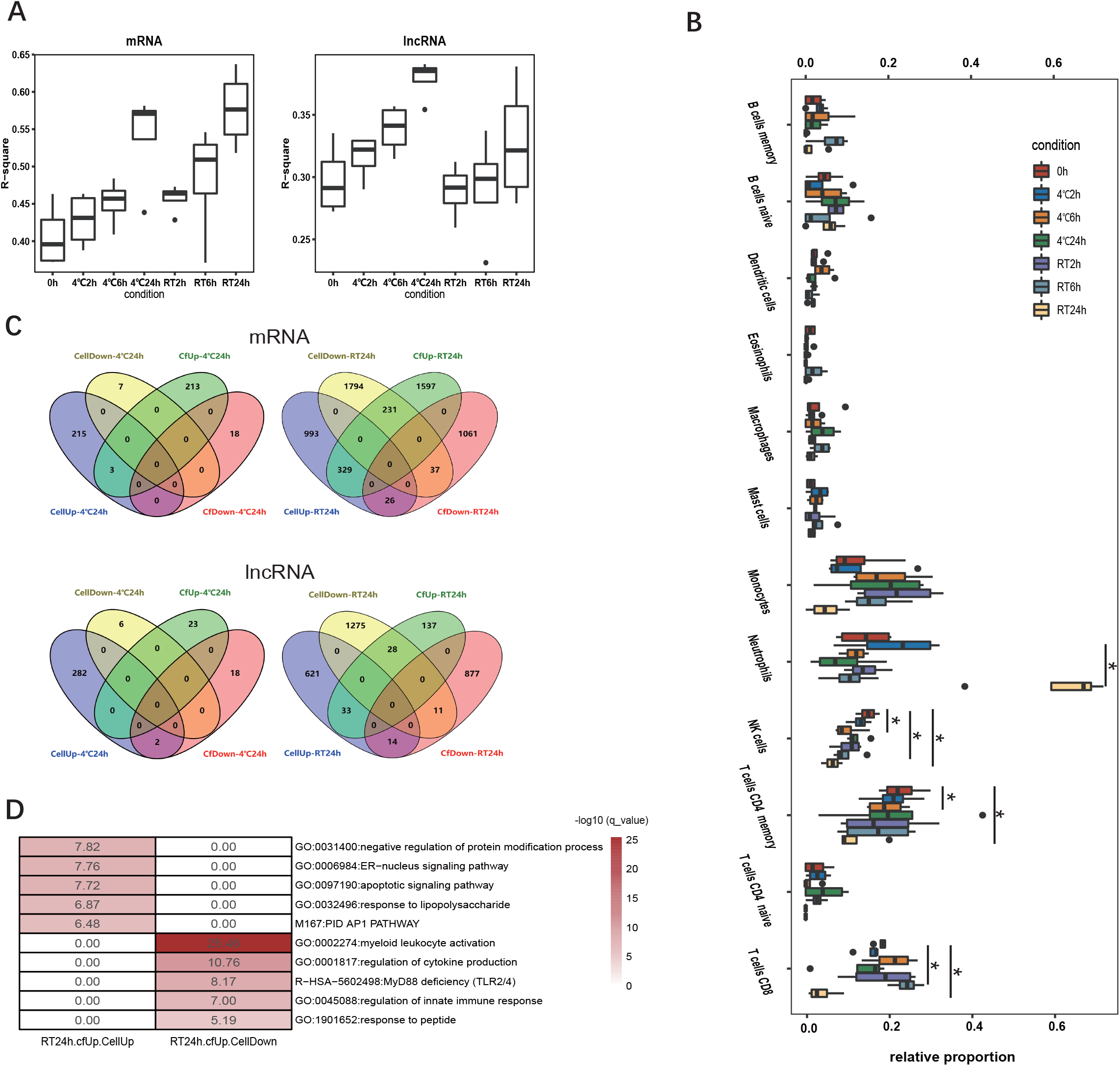
Effect of leukocyte on plasma cfRNA during blood storage (A) Correlations of gene expression between plasma cfRNA and leukocyte RNA. R^2^ is the square of Pearson’s correlation coefficient. (B) The overlap DEGs number of leukocyte RNA and plasma cfRNA at 24h, the DEGs of leukocyte RNA were listed in Supplementary Table 2. (C) Enriched pathway of leukocyte and plasma overlap DEGs. (D) leukocyte relative proportions of plasma cfRNA, estimated by deconvolution algorithms. * *p* < 0.05

To gain further insights into the impact of leukocyte on plasma RNA profile during blood storage, we performed cell type deconvolution analysis on cf-mRNA data to estimate the leukocyte’s relative proportion in plasma. Deconvolution result showed that the three largest proportions of plasma cf-mRNA were CD4 memory T cells (23.1%), CD8 T cells (18.4%) and neutrophils (14.3%) on average at 0h (Fig.4B, Supplementary Table 4), which is different from blood, in which neutrophils are the most plentiful cell type. We next compared the relative abundances of member cell types in plasma cfRNA under different storage conditions. When blood was stored at 4◻, there were little variations across time points, except the decreasing proportions of natural killer [25] cells and CD4 memory T cells at 6h (Fig.4B, Supplementary Table 4). When blood was stored at RT, the contribution of neutrophils sharply increased to 61.2% (*p* = 0.004) at 24h (Fig.4B, Supplementary Table 4), suggesting neutrophils RNA release increases after blood storage for 24h at RT. Meanwhile, relative fractions of CD8 T cells (*p* = 0.003), CD4 memory T cells (*p* = 0.004) and NK cells (*p* = 0.023) significantly decreases at 24h, which probably due to the increase of neutrophils.

To test whether the changes of plasma RNA profile are related to the changes in leukocytes during blood storage, we evaluated the number of DEGs that overlaped between leukocyte and plasma under the same storage conditions (Supplementary Table 1, Supplementary Table 5). As a result, there were few genes common between leukocyte and plasma RNA at 4◻ for 24h or at RT for 6h (Fig. 4C), maybe because of the small number of DEG. However, when blood was stored at RT for 24h, there were 329 up-regulated mRNA genes and 33 lncRNA genes of cfRNA overlapped with leukocyte up-regulated genes, which is significantly (*p* = 0.001) more than randomly permuting overlap gene number (Fig. 4C, Supplementary Fig. 3). The overlapped up-regulated genes were mainly enriched in the pathway of negative regulation of protein modification process, ER-nucleus signaling pathway, apoptosis, response to lipopolysaccharide and PID AP1 (Fig. 4D). At the same time, there were 231 plasma up-regulated coding genes overlap with leukocyte down-regulated genes (Fig. 4C), and these genes are mainly enriched in the pathway of myeloid leukocyte activation, regulation of cytokine production, MyD88 deficiency, regulation of innate immune response, and response to peptide (Fig. 4D). We noticed that the neutrophil activation and degranulation pathway is a member of the myeloid leukocyte activation pathway (Supplementary Table 6).

### Effect of blood storage on tissue enriched genes

A potential application of plasma cfRNA is to dynamically monitor tissue health or disease status by using tissue-specific genes, whose expression may be impacted by the blood storage conditions and lead to an unreliable result. In this study, TEGs of 23 tissues were taken into to evaluate, and only tongue enriched genes didn’t detect in the plasma. The highest detection ratios of TEGs are lymphoid tissue and bone marrow, followed by liver, with detection rates of 59.3%, 58.6%, and 32.2%, respectively (Fig. 5A).

**Fig. 5.**
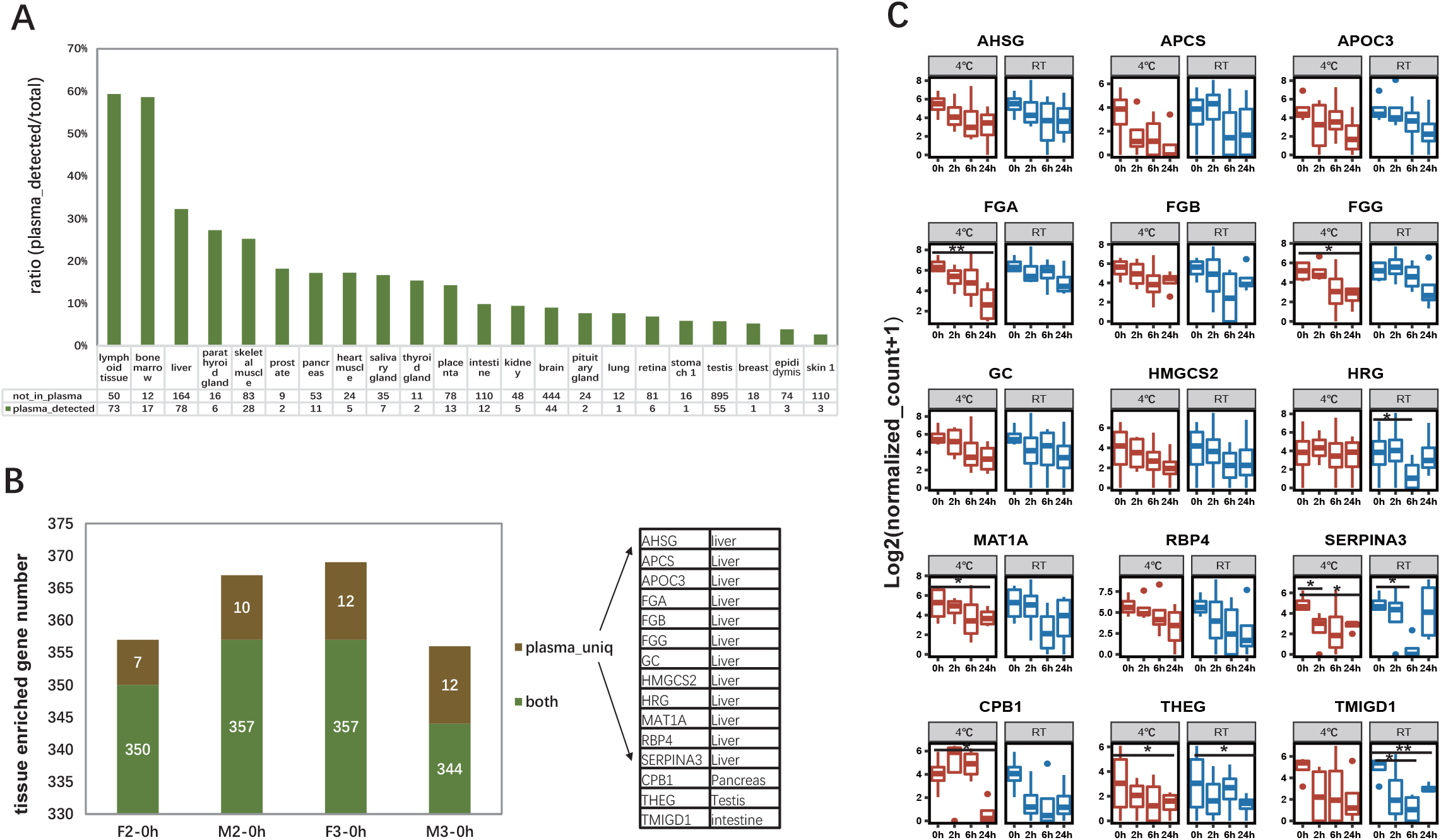
Effect of blood storage on tissue enriched genes (A) Gene detected ratios of TEGs in plasma cfRNA. (B) TEGs detected only in plasma are mainly liver-specific. (C) Dynamic expressions of plasma unique TEGs. The raw counts were normalized by DESeq2. ** different expression genes, that is log2(fold change) ≥ 1 and adjusted p-value <0.05; * genes with moderately changed expression, that is only *p* < 0.05.

To characterize the effects of blood storage on primary cfRNA, which is released from other tissues and can’t be supplemented by leukocytes during blood storage, we removed all genes that were detected in leukocytes at any conditions. Most TEGs (> 96%) were removed, and only 15 genes remained, among which 12 genes are liver-specific, one gene is pancreas-specific, one gene is testis-specific, and one gene is intestine-specific (Fig. 5B). The expressions of nearly all remaining TEGs were decreasing gradually, regardless of at 4◻ or at RT. Moreover, FGA and TMIGD1 were significantly down-regulated with adjusted p-value < 0.05 at 4◻ and RT after 24h incubation, respectively (Fig. 5C).

## Discussion

Plasma cfRNA is considered as a potential biomarker for disease diagnosis and monitoring [31]. However, many factors are known to dynamically influence the stability of cfRNA profile, including pre-analysis variables, such as blood storage time and temperature. Here, we characterized the effects of storage conditions on plasma cf-mRNA, cf-lncRNA and cf-miRNA, and integrated leukocyte RNA to further investigate the contamination of cell RNA and the decay of the original RNA.

To the best of our knowledge, this is the first study to comprehensively analyze the effect of blood storage conditions on plasma whole transcriptome profile. Most relevant studies only focused on specific genes [32, 33]. By evaluating the variabilities of specific miRNA genes, such as hsa-miR-16-5p, these studies thought that there is minimal effect on the level of cf-miRNA when analyzed within 24 hours of collection in EDTA tube type at RT [32, 33]. However, our result showed that although hsa-miR-16-5p kept relatively stable at RT for 24 hours, the effect on plasma cf-miRNA was observed by PCA and DEG analysis. Aoife Ward Gahlawat et al. also reported that the breast cancer-associated miRNA, namely miR-148b and miR-652 were significantly up-regulated after storage for 18 hours at RT [6]. A similar situation occurred in plasma mRNA. Previous studies indicated that plasma free mRNA is relatively stable at 4◻ for 24 hours [34] or at RT for 3 hours [35] by evaluating the influence of blood storage conditions from the expression of specific genes, such as housekeeping genes ACTB and GAPDH. In our study, although ACTB and GAPDH were relatively stable at 4◻ for 24 hours, 1.39% of the mRNA genes had changed significantly. The delayed processing of blood samples is inevitable in daily clinical practice sometimes, therefore considering the changes in the whole expression profile, we recommend that blood processing should be within 6 hours at 4°C and within 2h at RT to avoid confounding variables influencing the mRNA and lncRNA expression profile. For miRNA, we recommend that blood can be stored at 4◻ for 24h but should process within 6 hours at RT to minimize effects on cf-miRNA expression. In addition, the genes that are sensitive to blood storage conditions need to be interpreted carefully in future studies.

Our results showed a decreasing trend regarding the correlations of cfRNA expression between 0h and other storage conditions, while the correlations of plasma RNA and leukocyte RNA exhibited an increasing trend over time. We speculated that this phenomenon may result due to the contamination of leukocyte and platelet RNA as well as the degradation of plasma original RNA during the processing and storage period. Several studies had shown that plasma RNA concentrations increases over time when plasma is incubated with whole blood at RT, suggesting the RNA supplement from other components of blood [6, 36]. Our study shows that the average proportion of neutrophils in plasma grows at RT after 24h, while the proportion of neutrophils in blood cells decrease with storage time at RT [37, 38], suggesting an increase of neutrophils RNA contamination, which may be because of the short half-life of neutrophils. Also, the plasma up-regulated genes that overlapped with leukocyte down-regulated genes were enriched in the pathway of neutrophils activation and degranulation, which may be because that the neutrophil-specific genes are also the genes of neutrophil activation and degranulation pathway. All of these imply that neutrophils RNA is an important source for cellular RNA contamination for plasma cfRNA. Another important source of RNA contamination may be platelet [39]. Platelet-activation pathways were observed in plasma up-regulated mRNA genes at RT after 24h incubation of whole blood, and activated platelet can produce microparticles circulating in blood [40], which may increase platelet-specific cfRNA in the plasma.

Due to the influence of storage conditions, leukocyte may undergo some events such as apoptosis and necrosis that cause the cfRNA released, resulting in increased expression of specific genes in the plasma. We found that the overlapped genes of plasma and leukocyte up-regulated genes are enriched in the process of negative regulation of protein modification and endoplasmic-reticulum-nucleus signaling, which is activated by accumulation of unfolded proteins and may induce cell apoptosis [41]. Besides, the apoptosis signaling pathway is noticed in both up-regulated genes, suggesting that leukocyte is undergoing apoptosis and these cfRNA can also be released to plasma.

cfRNA is stable and protected by some mechanisms as more than 99% of the free added RNA was degraded after a 15-s incubation time but no degradation was observed in plasma cfRNA [32, 34]. In our study, we found that plasma down-regulated genes of 24h at RT were enriched in ribosomal protein mRNA, indicating that ribosomal protein mRNA may have different protection mechanisms from other genes, which needs more evidence. Besides, the expression of TEGs that was only detected in plasma has a trend of decreasing over time, indicating the degradation of plasma original mRNA. This suggests that the declination of plasma original RNA such as tissue-specific RNA, which is important when using plasma cfRNA to monitor the health and disease status of other tissues.

Future studies involving a large sample size along with leukocyte miRNA data is anticipated to provide further insights on the effect of leukocyte on plasma miRNA.

## Conclusions

In summary, by evaluating the changes of the whole transcriptome profile, we concluded that plasma cf-miRNA is relatively stable at 4◻ for 24 hours and at RT for 6 hours, while cf-mRNA and cf-lncRNA are relatively stable at 4◻ for 6 hours and at RT for 2 hours. Besides, the contamination of leukocyte and platelet RNA increased dramatically after 24 hours incubation of whole blood at RT. Furthermore, the decay of tissue-specific genes was continued over time, regardless of 4◻ or RT. This study highlights the importance of proper sample storage and facilitates further researches or applications to avoid biased results during sample processing.

## Author Contributions

Project administration, W-J.W.; Sample collection, S.Z. and J.Z.; data analysis, J.S. and T.W.; Methodology, X.Y., Y.X., Q.Z. and H.C.; Writing and review, W-J.W. and J.S.; Supervision, W-J.W., X.Z., and F.C.

## Acknowledgments

We would like to thank all participants for supporting this study, and appreciate that Sunil Kumar Sahu helps us modify the manuscript in grammar. We also sincerely thank the support provided by China National GeneBank.

